# Effects of body mass index on relationship status, social contact, and socioeconomic position: Mendelian Randomization study in UK Biobank

**DOI:** 10.1101/524488

**Authors:** Laura D Howe, Roshni Kanayalal, Robin N Beaumont, Alisha R Davies, Timothy M Frayling, Sean Harrison, Samuel E Jones, Franco Sassi, Andrew R Wood, Jessica Tyrrell

**Author notes:** Corresponding author – Oakfield House, Oakfield Grove, Bristol BS8 2BN. Contributor statement: LDH and JT conceived the study. Statistical analyses were performed by JT and RK. LDH, RK and JT wrote the first draft of the manuscript. All authors contributed to study design, interpretation of results and provided comments and critical revisions for intellectual content. LDH and JT accept full responsibility for the overall content as guarantors. The corresponding author attests that all listed authors meet authorship criteria and that no others meeting the criteria have been omitted. The Corresponding Author has the right to grant on behalf of all authors and does grant on behalf of all authors, a worldwide licence to the Publishers and its licensees in perpetuity, in all forms, formats and media (whether known now or created in the future), to i) publish, reproduce, distribute, display and store the Contribution, ii) translate the Contribution into other languages, create adaptations, reprints, include within collections and create summaries, extracts and/or, abstracts of the Contribution, iii) create any other derivative work(s) based on the Contribution, iv) to exploit all subsidiary rights in the Contribution, v) the inclusion of electronic links from the Contribution to third party material where-ever it may be located; and, vi) licence any third party to do any or all of the above. **What is known?** - Studies have demonstrated stigma and discrimination against people who are overweight or obese in social, educational and employment settings. - Mendelian Randomization, a technique that uses genetic data to overcome confounding and reverse causality, has provided evidence for a causal effect of higher BMI on lower socioeconomic position (SEP). It is reasonable to hypothesise that adverse socioeconomic effects might be particularly strong for very high BMI, and also evident for low BMI, but this has not been explored by previous studies. - It is not known whether BMI also has a causal effect on social outcomes, such as cohabitation with a partner/spouse, contact with friends/family, and participation in leisure activities. **What this study adds** - Using Mendelian Randomization, we found evidence of sex-specific effects of BMI on likelihood of being in a cohabiting relationship with a partner or spouse: in men, lower BMI was associated with being less likely to live with a partner or spouse, whereas in women, higher BMI was associated with being less likely to live with a partner or spouse. - We found evidence of causal effects of BMI on several domains of SEP (income, deprivation, education, skilled employment), with BMIs at both ends of the distribution (both high and low BMI) leading to lower income and higher levels of deprivation. Higher BMI was associated with lower participation in leisure activities in men but not women; effects of BMI on other measures of social contact were not observed.

## Abstract

**Objective:** To assess whether body mass index (BMI) has a causal effect on social and socioeconomic factors, including whether both high and low BMI can be detrimental.

**Design:** Mendelian Randomization, using genetic variants for BMI to obtain unconfounded estimates, and non-linear Mendelian Randomization.

**Setting:** UK Biobank.

**Participants:** 378,244 men and women of European ancestry, mean age 57 (SD 8 years).

**Main outcome measures:** Townsend deprivation index, income, age completed full time education, degree level education, job class, employment status, cohabiting relationship status, participation in leisure and social activities, visits from friends and family, and having someone to confide in.

**Results:** Higher BMI was causally associated with higher deprivation, lower income, fewer years of education, lower odds of degree-level education and skilled employment. For example, a 1 SD higher genetically-determined BMI (4.8kg/m^2^ in UK Biobank) was associated with £1,660 less income per annum [95%CI: £950, £2,380]. Non-linear Mendelian Randomization provided evidence that both low BMI (bottom decile, <22kg/m^2^) and high BMI (top seven deciles, >24.6kg/m^2^) can increase deprivation and reduce income. In men only, higher BMI was related to lower participation in leisure and social activities. There was no evidence of causal effects of BMI on visits from friends and family or in having someone to confide in. Non-linear Mendelian Randomization analysis showed that low BMI (bottom three deciles, <23.5kg/m^2^) reduces the odds of cohabiting with a partner or spouse for men, whereas high BMI (top two deciles, >30.7kg/m^2^) reduces the odds of cohabitation with a partner or spouse for women.

**Conclusions:** BMI affects social and socioeconomic outcomes, with both high and low BMI being detrimental for some measures of SEP. This suggests that in addition to health benefits, maintaining healthy ranges of BMI across the population could have benefits both for individuals and society.

## Introduction

Lower socioeconomic position (SEP) is associated with higher body mass index (BMI) and greater risk of obesity in both children and adults, with stronger social patterning of obesity seen in women compared with men.^1–4^ Social factors such as relationship status and social contact networks are also related to BMI and other aspects of health.^5–7^ Higher BMI is associated with stigma, discrimination, lower self-esteem, and physical and mental ill-health^8–15^, all of which could potentially affect social, educational, and employment outcomes, meaning that the relationships between SEP, social factors and BMI could be bidirectional. A study conducted by Flint et al.^16^ demonstrated considerable weight bias in the recruitment process; employers across various sectors rated the same curriculum vitae as less suitable for a job when it was accompanied by a photograph of a person with obesity, particularly a woman with obesity. Actions to address the increasing prevalence of obesity across the UK are supported by the strong links between BMI and health outcomes, but demonstrating effects of BMI on social and socioeconomic outcomes could augment the impetus for policy makers across sectors to act to prevent obesity, with the potential for greater societal benefits.

A key challenge is the strong influence of social factors on BMI, which means that when we come to study the downstream social and socioeconomic consequences of BMI, earlier life social factors can lead to confounding and reverse causality bias. Natural experiments are therefore needed to obtain unbiased estimates of the effects of BMI on social and socioeconomic outcomes. One example of a natural experiment is sibling comparisons; siblings share their family environment and therefore if siblings discordant for obesity also differ with respect to SEP, this suggests the association is not due to confounding by shared family-level factors. In a study of male Swedish siblings, men who were obese as teenagers (n=2,600, of whom 95% had a sibling who was not obese) had an income 9% lower than men who were not obese as teenagers.^17^ This sibling-comparison design provided evidence that both high and low BMI are associated with lower SEP; underweight teenagers also had lower income than their normal BMI siblings.

Mendelian Randomization (MR) is an alternative natural experiment approach that uses genetic variants related to an exposure of interest (here, BMI) as instrumental variables.^18^ The approach exploits the natural experiment of genetic variants being randomly assigned at conception, which means that they are less likely to be associated with factors that would confound a traditional analysis and should not suffer from reverse causality. A previous MR study in 120,000 European UK Biobank participants provided evidence of a causal effect of higher BMI on several socioeconomic outcomes, particularly lower income and higher area-level deprivation in women.^19^ Using data from over 350,000 participants in UK Biobank, we build on this study by extending the analysis to additionally examine effects of BMI on important social outcomes - cohabiting relationship status and three measures of social contact. Given evidence that both low BMI and high BMI are associated with various adverse outcomes^17 20–22^, we use novel MR methods to assess non-linearities in the effects of BMI on social and socioeconomic outcomes.

## Methods

### UK Biobank

The UK Biobank is a study of over 500,000 individuals aged between 37 and 73 years (99.5% between 40 and 69 years) recruited between 2006 and 2010. The study is described in detail elsewhere ^23^ Participants attending one of 22 baseline assessment centres across the UK provided detailed information via questionnaires, interviews and measurements. Field codes for variables used are in the Supplementary Text.

### Patient Involvement

The current research was not informed by patient and public involvement because it used secondary data. This study utilised the UK Biobank resource, which has details on patient and public involvement available online (http://www.ukbiobank.ac.uk/about-biobank-uk/ and https://www.ukbiobank.ac.u/wp-content/uploads/2011/07/Summary-EGF-consultation.pdf?phpMyAdmin=trmKQlYdjjnQIgJ%2CfAzikMhEnx6).

### Exposure and outcome measures

The exposure and outcome measures were defined from the participants’ baseline visit to the UK Biobank assessment centre.

#### Body mass index

BMI was calculated from measured weight (kg)/height (m)^2^ and inverse normalised prior to analysis.

#### Socioeconomic position

We used six measures of SEP; full details in the Supplementary Text:

- Townsend deprivation index (TDI); an area-based measure at census-tract level; higher scores indicate higher deprivation.
- Annual household income.
- Job class; coded as skilled versus unskilled.
- Employment status; coded as employed (or self-employed) versus unemployed.
- Years in education.
- Degree status; coded as degree-level education or lower.

#### Cohabiting relationship status and social contact

Whether or not a participant was currently cohabiting with a partner or spouse was defined by their responses to a question about their relationship to other household members; participants reporting that they lived with a ‘husband, wife or partner’ were defined as being in a current cohabiting relationship. Participants also reported whether in the last two years they had experienced the ‘death of a spouse or partner’ or ‘marital separation/divorce’; these data did not form part of our definition of current cohabitation, but were used in sensitivity analysis.

We used three measures of social contact; full details in the Supplementary Text:

- Visits from friends and family: less than weekly versus weekly or more visits.
- Participation in leisure and social activity: any activity versus none.
- Confiding in others: less than weekly versus weekly or more.

### Sample definition

Our analyses are restricted to unrelated participants of white European ancestry, defined by Principal Component Analysis of genetic data (see Supplementary Text for full details). Participants were also excluded if they had missing data for BMI (N=1,494), or missing data for a given outcome (Ns varied by outcome, see Table 1).

**Table 1:** Characteristics of the 378,274 participants of European ancestry with valid genetic, BMI and at least one outcome measure

| Characteristic | All (n=378,244) | Men (n= 174,358) | Women (n=203,886) | P value* |
| --- | --- | --- | --- | --- |
| Mean age at recruitment in years (SD) | 57.2 (8.0) | 57.5 (8.1) | 57.0 (7.9) | <1x10<sup>-15</sup> |
| N male sex (%) | 174,358 (46.1%) | NA | NA | NA |
| Mean body mass index in kg/m<sup>2</sup> (SD) | 27.4 (4.8) | 27.8 (4.2) | 27.0 (5.2) | <1x10<sup>-15</sup> |
| **Socioeconomic position measures** |  |  |  |  |
| Mean Townsend Deprivation Index (SD) | -1.48 (2.99) | -1.44 (3.05) | -1.51 (2.94) | <1x10<sup>-15</sup> |
| Annual Household Income, N(%): |  |  |  |  |
| < £18,000 | 70,841 (18.7) | 30,536 (17.5) | 40,305 (19.8) | <1x10<sup>-15</sup> |
| £18,000 to £30,999 | 82,818 (21.9) | 38,224 (21.9) | 44,594 (21.9) |  |
| £31,000 to £51,999 | 86,024 (22.7) | 42,443 (24.4) | 43,581 (21.4) |  |
| £52,000 to £100,000 | 68,240 (18.0) | 35,598 (20.4) | 32,642 (16.0) |  |
| > £100,000 | 18,194 (4.8) | 9,720 (5.6) | 8,474 (4.2) |  |
| Mean number of years spent in education (SD) | 15.0 (5.1) | 15.3 (5.1) | 14.6 (5.1) | <1x10<sup>-15</sup> |
| Number with a degree, N(%) | 179,444 ( 47.4) | 83,869 (48.1) | 95,575 (46.9) | <1x10<sup>-15</sup> |
| Number with a skilled job, N(%)** | 199,992 (81.6) | 97,908 (83.1) | 102,084 (80.2) | <1x10<sup>-15</sup> |
| Number of participants in employment, N(%) | 216,248 (57.2) | 104,979 (60.2) | 111,269 (54.6) | <1x10<sup>-15</sup> |
| **Social contact measures** |  |  |  |  |
| Cohabit with a partner or spouse, N(%) | 277,399 (73.3) | 134,062 (76.9) | 143,337 (70.3) | <1x10<sup>-15</sup> |
| At least weekly visits from friends or family, N(%) | 294,864 (78.0) | 127,627 (73.2) | 167,237 (82.0) | <1x10<sup>-15</sup> |
| Weekly participation in leisure and social activity, N(%) | 262,868 (69.5) | 121,596 (69.7) | 141,272 (69.3) | 0.031 |
| At least weekly opportunities to confide in someone, N(%) | 274,092 (72.5) | 119,322 (68.4) | 154,770 (75.9) | <1x10<sup>-15</sup> |

\*P values were determined in age and sex adjusted linear and logistic regression models \*\*Percentage given as those from those reporting a job class

### Statistical analysis

For TDI, years in education and income, we converted the data to a normal distribution to limit the influence of any subtle population stratification and to provide standard deviation (SD) effect sizes. Analyses using these normalised variables were then adjusted for eight covariates: age, sex, assessment centre location and five (within-European) ancestry principal components. To convert our results back to meaningful units after analysis, we multiplied our SD betas by the SD of the outcome. For example, the SD of TDI was 3.0 units. Therefore, a difference of 0.05 SD equated to a 0.15 unit difference in deprivation. Analyses using the original variables within ordinal logistic regression models yielded the same pattern of results and are therefore not described further.

#### Linear and logistic regression

We regressed each outcome measure against BMI using linear regression for continuous outcome variables and logistic regression for binary outcomes. Age and sex were included as covariates.

#### Genetic risk score for BMI

Genetic variants for BMI were selected from UK Biobank’s imputation dataset.^24^ We selected 73 of 76 common genetic variants associated with BMI at genome-wide significance in the GIANT consortium studies of up to 339,224 people (Supplementary Table 1).^25^ The variants selected were limited to those associated with BMI in the analysis of all people of European ancestry and did not include those that reached genome-wide levels of statistical confidence in only one sex or one stratum. Variants were also excluded if they were known to be classified as a secondary signal within a locus: three variants were excluded from the score because they were known to have pleiotropic effects on other traits (rs11030104 (*BDNF* reward phenotypes including smoking), rs13107325 (*SLC39A8* lipids, blood pressure), rs3888190 (*SH2B1* multiple traits)).^19 25^

The 73 BMI variants were combined into a weighted genetic risk score (GRS). Each variant was weighted by its effect size (β-coefficient) obtained from the primary GWAS that did not include any UK Biobank data (Equation 1).^25^ The weighted score was then rescaled to reflect the number of trait-raising alleles (Equation 2).

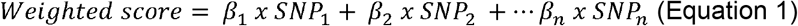

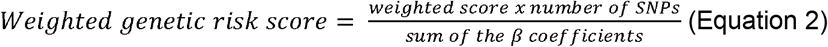

#### Mendelian randomisation

MR was utilised to investigate causal pathways from BMI to SEP, cohabitation, and social contact. We employed the two-stage-least-squares regression estimator method that uses predicted levels of BMI per genotype and regresses the outcome against these predicted values. For continuous outcomes we used the ivreg2 command in Stata. For binary outcomes, the analysis was done in two stages. Firstly, the association between the BMI GRS and BMI was assessed. The predicted values from this regression were used as the independent variable (reflecting an unconfounded estimate of variation in BMI) and the binary or ordinal outcomes as the dependent variable in logistic regression models. Robust standard errors were used.

#### Differences between men and women

To test the hypothesis that the effects of BMI on SEP, cohabitation and social contact may differ in men and women, we repeated linear and logistic regression and MR analyses separately in each sex. The selected BMI genetic variants have very similar effects in men and women and therefore the same variants and GRS were used in all participants and in sex-stratified analyses. The beta values for men and women were compared using Fisher’s z-score method (equation 3).^26^

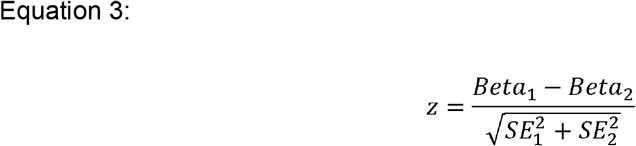

#### Sensitivity Analyses

##### i. Mechanisms underlying associations between BMI and cohabitation

Any associations between BMI and cohabitation with a partner/spouse could result from i) associations between BMI and partnership formation, ii) associations between BMI and separation/divorce, iii) mortality of partners – if people are likely to form relationships with someone whose BMI is similar to theirs, this could result in people of high BMI being more likely to have partners who also have high BMI, and hence who are more likely to die at a younger age. To help unpick these mechanisms, we examined the associations between BMI and i) separation from partner/spouse in the past two years, and ii) death of a partner or spouse in the past two years. Due to low numbers of people reporting these two events, these analyses were limited to logistic regression; no MR was performed. Data on lifetime history of cohabiting relationships is not available in UK Biobank. We also repeated our main logistic regression and MR analyses for cohabitation stratifying by median age, <58 years or 58 or more years, with the hypothesis that if partner death was contributing to the overall associations, the association would be stronger in the older age group where mortality rates are higher.

##### ii. Effect of household size on associations of BMI with income

We explored the causal relationships between BMI and annual household income per capita by adjusting the income variable for the number of people living in the household. It is not possible with UK Biobank data to ascertain how many of the people in a household are adults and children.

##### iii. Understanding whether associations are independent of ill health

We repeated our analyses in a subset of 88,323 individuals with no known health problems to investigate whether associations were driven by poor health.

##### iv. Evaluating pleiotropy: negative control, FTO as an instrument, and two-sample MR

A potential source of residual confounding in MR studies is horizontal pleiotropy.^27^ As a negative control, we used logistic regression and MR to assess the association between BMI and use of sun protection (based on self-report of whether the participant ‘usually or always’ versus ‘never or sometimes’ uses sun protection (e.g. sunscreen lotion, hat) when outdoors in summer. Lower SEP is associated with lower use of sun protection, but BMI is unlikely to have a causal effect on sun protection use other than a path from BMI to SEP to sun protection use, which means associations between BMI and sun protection use should be much weaker than those seen for associations of BMI with SEP. As another way of excluding pleiotropic effects of the BMI GRS, we repeated our analysis using only FTO (rs1558902), the genetic variant with the largest effect on BMI. This analysis is likely to have low power, but is unlikely to exhibit pleiotropy. A range of two-sample MR analyses were performed for linear MR models, to assess the validity of the MR assumptions.^27^ Full details of the methods used^28^ ^29^ are in Supplementary Text.

##### v. Using a larger BMI genetic risk score

Recently, further genetic variants for BMI have been identified using approximately 700,000 individuals of European ancestry^30^, including individuals from the UK Biobank. These newly discovered 941 variants explain more variation in BMI (6% compared with <3% for the GIANT study^25^) and hence represent a more powerful tool for MR. We did not use a GRS based on this study as our main analysis because they were discovered using the UK Biobank and therefore may bias our findings. However, as a sensitivity analysis, we repeat our MR analysis using this GRS.

#### Analysis of non-linearity

To explore potential non-linear relationships between BMI and each outcome, we employed the nlmr package (https://github.com/jrs95/nlmr) in R^31^ to assess how the association of BMI with each outcome differs across deciles of IV-free BMI (residuals from a regression of BMI on the genetic instrumental variable). Full details are in the Supplementary Text.

## Results

BMI was available for 378,244 unrelated individuals of white European ancestry with at least one of the outcome measures (Table 1). The 73 SNP BMI GRS was robustly associated with BMI, explaining 1.7% of the variance. The 941 SNP BMI GRS explained 5.0% of the variation in BMI.

### Association of BMI with socioeconomic position

In linear and logistic regression models, higher BMI was associated with higher deprivation, lower income, fewer years in education and lower odds of holding a degree, being employed, and working in a skilled profession (Figures 1A-1F; Supplementary Table 2).

**Figure 1:**
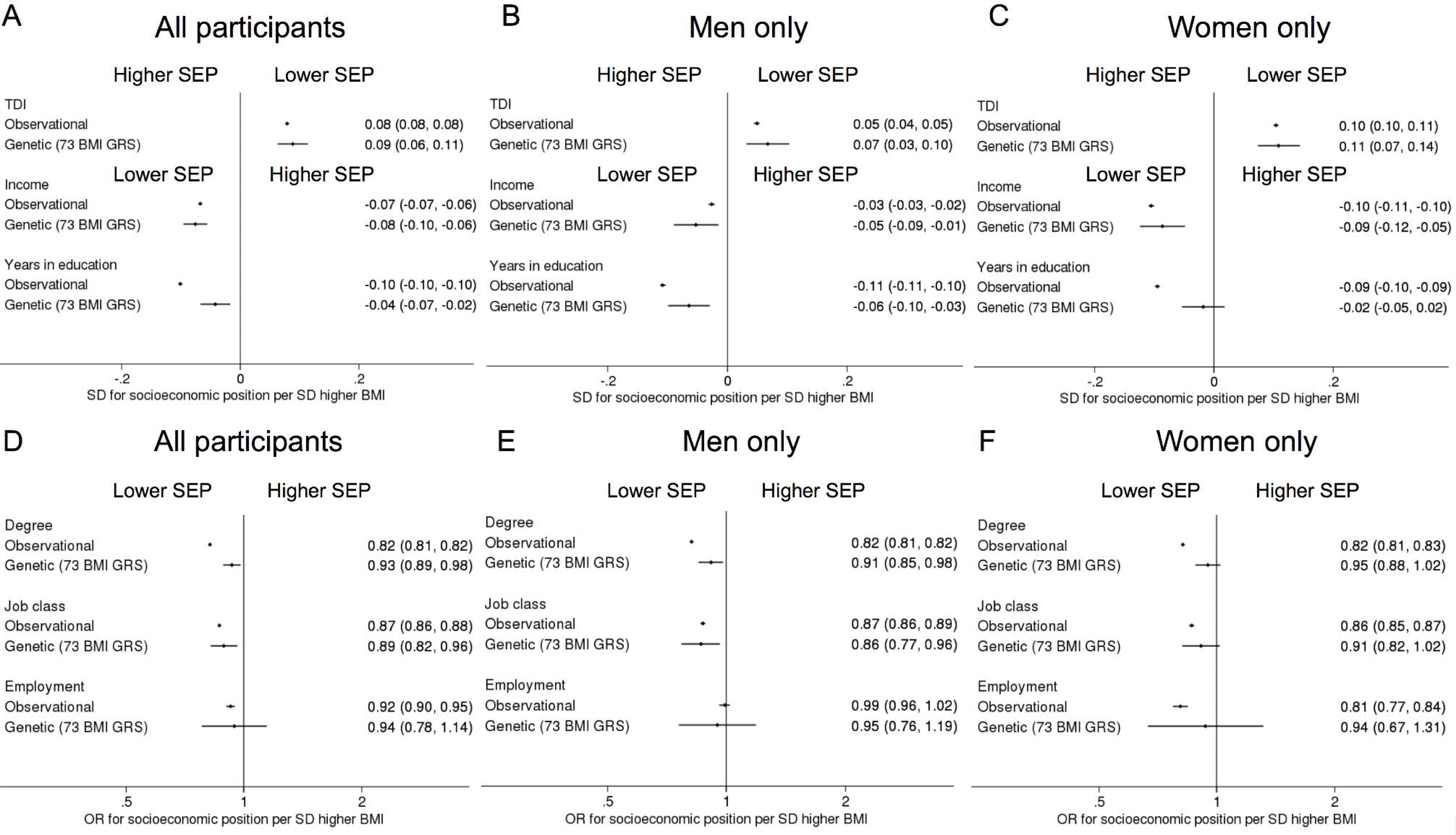
Forest plots of the linear and logistic regression and MR associations between a 1 SD higher BMI and 6 measures of socioeconomic position (SEP). Panels A-C represent the differences in continuous SEP measures (TDI, income and years in education) in A) all individuals, B) men only and C) women only. Panels D-F represent differences in binary SEP measures (degree, skilled job and employment status) in D) all individuals, E) men only and F) women only. Higher and lower SEP are marked on the plots as higher SEP and lower SEP. Note Townsend deprivation index (TDI) is the opposite way round to the other SEP measures, with higher values representing more deprivation.

Mendelian Randomization provided evidence for a causal role of higher BMI on lower SEP (Figures 1A-1F; Supplementary Table 2). In men and women combined, a 1 SD higher genetically-determined BMI (4.8kg/m^2^ in UK Biobank) was associated with a) 0.09 SD higher deprivation [95%CI: 0.06, 0.11, p=4×10^−12^], which equates to 0.27 unit higher level of deprivation, or approximately one third of a decile, b) 0.07 SD lower income [95%CI: 0.04, 0.10, p=4×10^−7^] which approximates to £1,660 less income per annum [95%CI: £950, £2,380], c) 0.04 SD fewer years of education [95%CI: 0.02, 0.07, p=0.001], which equates to approximately 3 months, d) 7% less chance of having a degree [odds ratio (OR) 0.93, 95%CI: 0.89, 0.98, p=0.007] and e) 11% less chance of having a skilled job [OR 0.89, 95%CI: 0.82, 0.96, p=0.003]. There was little evidence of a causal effect of BMI on being employed; OR=0.94, 95% CI 0.78, 1.14, p=0.6. Statistical tests for sex differences provided little evidence of associations between BMI and SEP differing between men and women (Supplementary Table 2).

### Association of BMI with relationship status

Higher BMI was associated with higher odds of cohabitation with a partner or spouse in men, and lower odds of cohabitation with a partner or spouse in women in logistic regression models (Figure 2; Supplementary Table 2). MR analysis provided evidence for a causal role of higher BMI on lower odds of cohabitation with a partner or spouse in women [OR: 0.83, 95%CI: 0.76, 0.92, p=0.0002]. In men, there was little evidence for an association between BMI and cohabiting with a spouse/partner [OR: 1.07, 95% CI: 0.97, 1.17, p=0.2]. There was strong evidence for sex differences in the MR results, p=0.0004.

**Figure 2:**
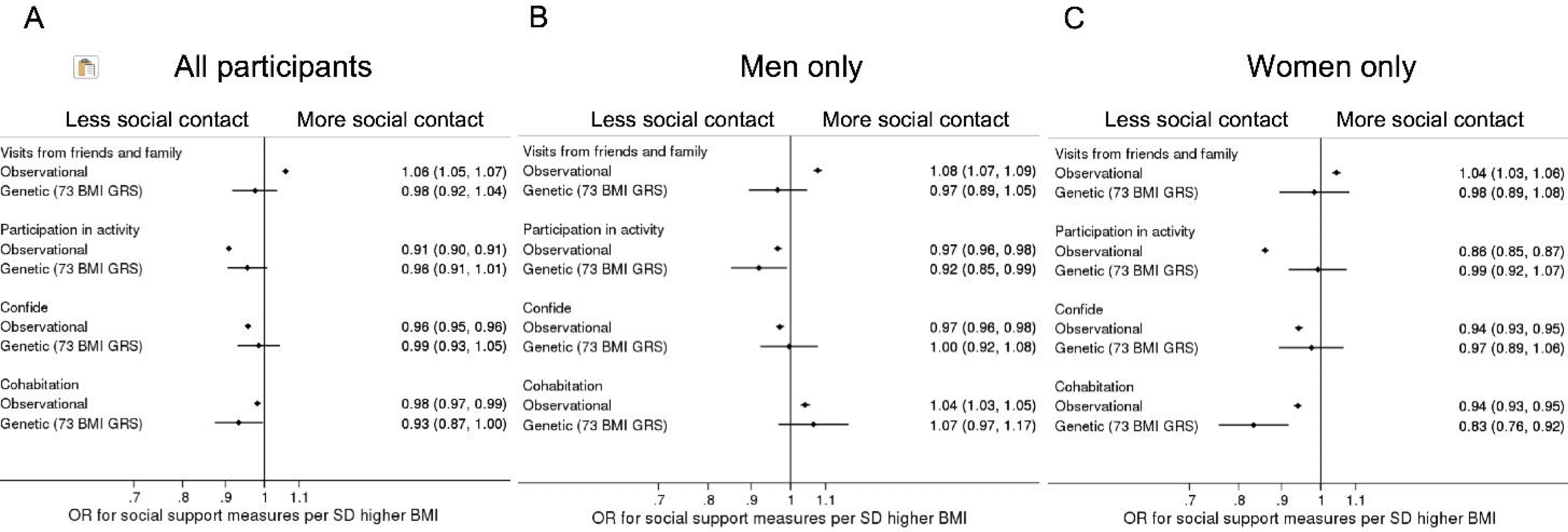
Forest plots of the linear and logistic regression and MR associations between a 1 SD higher BMI and the 4 social contact measures in A) all individuals, B) men only and C) women only. Higher and lower social contact is marked on the plots as more social contact and less social contact.

### Association of BMI with social contact

In logistic regression analyses, higher BMI was associated with higher odds of seeing friends and family on a weekly basis, lower odds of having someone to confide in regularly and lower odds of participating in leisure activities (Figure 2, Supplementary Table 2). In MR analyses, higher BMI was associated with lower odds of participation in leisure and social activities in men [OR: 0.92, 95% CI: 0.85, 0.99, p=0.03] but not women [OR: 0.99, 95% CI: 0.92, 1.07, p=0.9], although the test for sex differences did not provide strong evidence for interaction (p=0.2). There was no evidence for a casual role of BMI on the odds of weekly visits from friends and family or having someone to regularly confide in.

### Sensitivity analyses

#### i. Mechanisms underlying associations between BMI and cohabitation

Divorce or marital separation in the past two years was experienced by 7,745 participants, and 4,415 participants reported death of a partner or spouse in the last two years. Logistic regression analyses in men and women combined demonstrated that higher BMI was associated with lower odds of divorce/separation in the past two years (Supplementary Table 3). When analyses were stratified by gender, this association was seen in women [OR 0.95, 95% CI 0.92 to 0.97, p=2×10^−5^] but not in men [OR 1.02, 95% CI 0.99 to 1.05, p=0.14]. Higher BMI was associated with higher chance of a partner or spouse having died in the last two years in all participants [OR 1.05, 95% CI 1.02 to 1.07, p=0.001], and both sexes in stratified analyses. In age stratified analyses (Supplementary Table 4), MR results indicate a strong association between higher BMI and lower chance of being in a cohabiting relationship in people younger than 58 years, which is limited to women [OR in women 0.74, 95% CI 0.64 to 0.85, p=1×10^−5^; OR in men 1.07, 95% CI 0.94 to 1.21, p=0.3]. These associations were not seen in people aged 58 years and older [OR in women 0.93, 95% CI 0.81 to 1.06, p=0.3; OR in men 1.08, 95% CI 0.94 to 1.24, p=0.3].

#### ii. Effect of household size on associations of BMI with income

Accounting for the number of individuals in the household did not alter the relationship between BMI and income (Supplementary table 5).

#### iii. Understanding whether associations are independent of ill health

Linear and logistic regression and MR analyses were repeated in a subset of up to 88,223 individuals who reported no health problems; if the pattern of associations is similar in this ‘healthy’ subset of participants it would suggest health problems are not the (only) driver of associations between BMI and socioeconomic and social outcomes. Generally, the linear and logistic regression results were consistent with the main analyses (Supplementary table 6). In MR analyses, the associations were mostly in the same direction as the main analysis on the full cohort, but coefficients were generally closer to the null and confidence intervals were wider; there was only strong evidence for a causal role of higher BMI on higher deprivation in all individuals and women only (Supplementary table 6).

#### iv. Evaluating pleiotropy: negative control, FTO as an instrument, and two-sample MR

In logistic regression analyses, a one SD higher BMI was associated with a 6% lower chance of our negative control outcome, using regular sun protection [OR 0.94, 95% CI 0.94 to 0.95, p<1×10^−15^]. In MR analysis, the association was weak but still present (OR 0.98, 95% CI 0.97 to 0.99, p=0.002). Using FTO only, all associations had wide confidence intervals and large P values. Many (deprivation, income, years in education, participation in leisure and social activities in men), but not all (skilled job, employment) of the results were directionally consistent with our main analysis (Supplementary Table 7). There was some evidence of pleiotropy when years spent in education or having a degree were the outcome variable when using Egger-MR, but not for other outcomes. MR approaches using methods robust to pleiotropy provided findings that were largely consistent with our main analysis, with the greatest degree of consistency for the two continuous outcomes, income and deprivation, which have the highest statistical power (Supplementary Table 2).

#### v. Using a larger BMI genetic risk score

Using the 941 BMI variant GRS generally provided evidence consistent with our main analysis (Supplementary Table 2).

### Non-linear relationships

Using non-linear MR, we assessed the association of BMI with SEP, cohabitation status and social contact across deciles of ‘IV-free’ BMI (residuals from a regression of BMI on the BMI genetic risk score; hereon referred to as ‘BMI deciles’ for brevity). Full results are presented in Supplementary Tables 8 and 9. For deprivation, there was evidence that both low and high BMI resulted in higher deprivation (Figure 3). For people in the lowest BMI decile (<22kg/m^2^), higher BMI was causally associated with lower deprivation. Detrimental effects of high BMI on deprivation were evident for the top seven BMI deciles (>24.6kg/m2). The adverse effects of the lowest decile (<22kg/m^2^) were approximately equivalent to the adverse effects of BMI in the highest decile (>33.4kg/m^2^).

**Figure 3:**
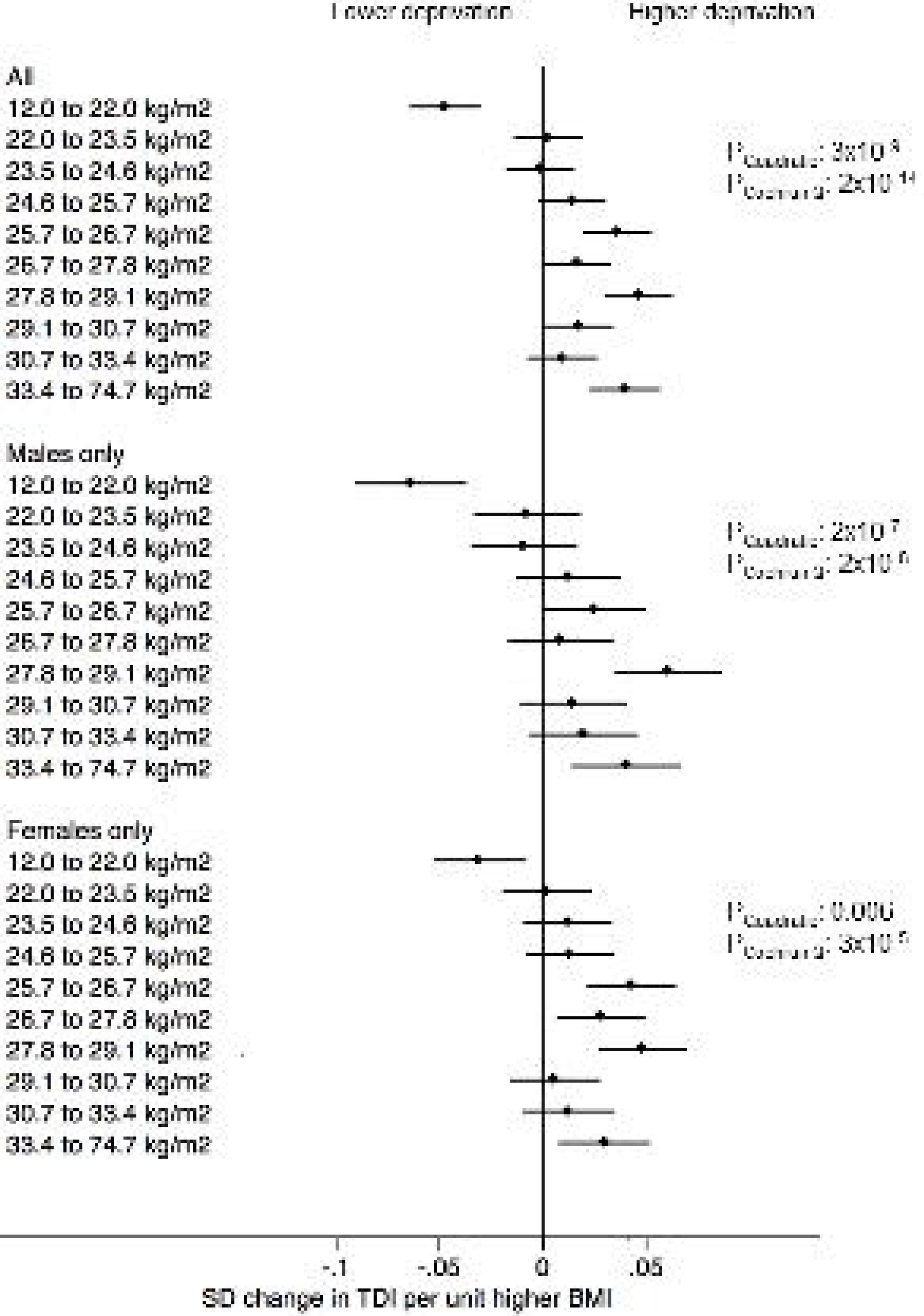
Dot plot exploring the non-linear association between BMI and Townsend deprivation index in deciles of genetically-determined BMI using the Piecewise Linear Model from non-linear MR analysis. The p-values presented for causal non-linear relationships are from the Piecewise Linear Model (P_Quadratic_ and P_Cochran Q_). Full results of non-linear analyses are provided in Supplementary tables 8 and 9.

Similar to the results for deprivation, both low and high BMI were also associated with lower income (Supplementary Figure 1). For people in the lowest two BMI deciles (<23.5kg/m^2^), higher BMI was causally associated with higher income. Detrimental effects of high BMI on income were evident for BMIs above 24.6kg/m^2^ (top seven tenths). The adverse effects of low BMI were of greater magnitude than the adverse effects of high BMI.

Sex differences were apparent in the non-linear MR results for cohabitation with a partner or spouse (Figure 4). In women, the results suggested an association between higher BMI and lower odds of cohabitation with a partner or spouse. This association was strong for the top two BMI deciles (equivalent to >30.7kg/m^2^). For example, in the top decile (>33.4kg/m^2^), each one SD higher BMI was associated with an 8% lower chance of cohabiting with a partner or spouse [OR 0.92, 95% CI 0.87 to 0.96, p=0.0002]. In contrast, in men there was strong evidence that low BMI (bottom three deciles, <24.6kg/m^2^) was associated with reduced odds of cohabitation with a partner or spouse, for example in the lowest decile (<22kg/m^2^), each one SD higher BMI was associated with 18% higher chance of cohabiting with a partner or a spouse [OR 1.18, 95% CI 1.13 to 1.24, p=1×10^−11^].

**Figure 4:**
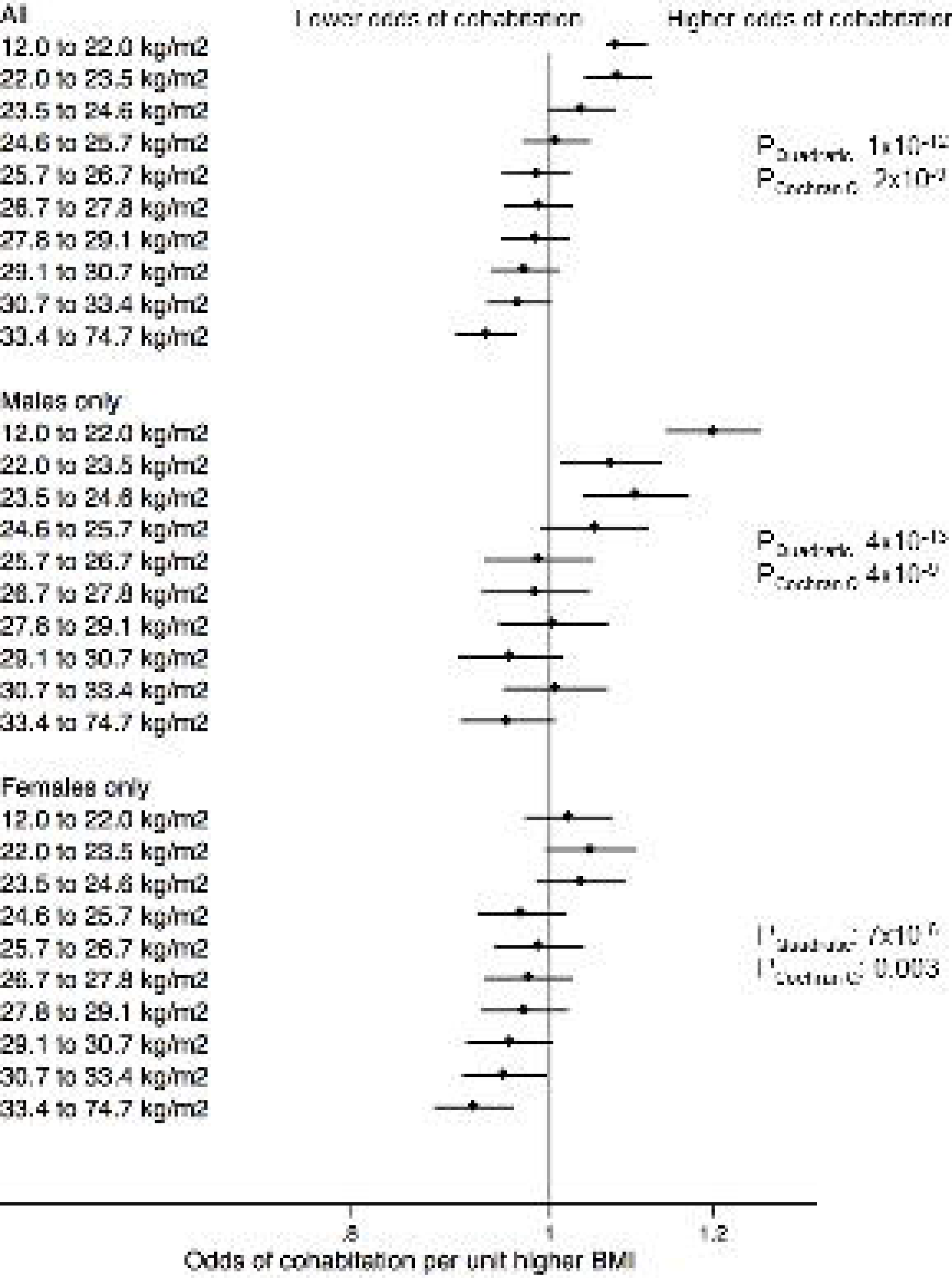
Dot plot exploring the non-linear association between BMI and the odds of cohabitation in deciles of genetically-determined BMI using the Piecewise Linear Model from non-linear MR. The p-values presented for causal non-linear relationships are from the Piecewise Linear Model (P_Quadratic_ and P_Cochran Q_). Full results of non-linear analyses in Supplementary tables 8 and 9.

For other outcomes, MR analyses did not provide evidence of non-linear relationships.

## Discussion

The application of genetic variants as an instrumental variable for BMI in this study has obtained estimates for the effect of BMI on a range of social and economic outcomes that are less affected by confounding or reverse causality than traditional analyses. Our study provides evidence that both low BMI (<22kg/m^2^) and high BMI (>24.6kg/m^2^) can lead to lower SEP in both men and women. We also demonstrated that higher BMI can lead to lower odds of participation in leisure activities for men. Finally, we showed that in men, lower BMI (<24.6kg/m^2^) is associated with lower odds of being in a cohabiting relationship with a partner or spouse, whereas in women, high BMI (>30.7kg/m^2^) is associated with lower odds of being in a cohabiting relationship with a partner or spouse. We found no evidence of a causal effect of BMI on contact with friends and family or having someone to confide in.

Associations between high BMI and lower SEP could arise due to the mental and physical health consequences of high BMI, and/or via social mechanisms including cultural norms and expectations of body size and related stigma and discrimination.^10^ We observed a similar pattern of associations in the subset of participants reporting no health conditions, suggesting that ill health is not the only mechanism driving the effects of BMI. Whilst our results suggest higher BMI leads to lower income and lower likelihood of being in a skilled occupation, there was no evidence of a causal effect of BMI on overall employment, suggesting that BMI affects skill level of job or career progression rather than entry into/continued participation in the labour market.

The adverse effects of low BMI were of similar magnitude to the adverse effects of high BMI, mirroring the results of a previous sibling comparison study^17^, although the effects of high BMI were seen across a wider range of BMIs and a larger number of people. Since low BMI is unlikely to be a cause of many health problems, and since we use MR to ensure results are not affected by reverse causality, the associations we see between low BMI and lower SEP are unlikely to be driven by health. Associations between low BMI and higher deprivation and lower income were observed in the bottom decile of BMI, <22kg/m^2^. The majority of people in this group have a BMI within the ‘ideal’ range (18.5 to 24.9). For the association between low BMI and lower odds of cohabitation in men, this association is apparent for BMIs below 24.6kg/m^2^, i.e. encompassing almost the entire range of ‘ideal’ BMI. The mechanisms driving the effects of low BMI remain uncertain and require further analysis. Associations of BMI with cohabitating relationship status could arise due to effects of BMI on partnership formation, divorce/separation, and/or partner death (if partnerships are more likely to form within couples of similar BMI). We were unable to assess the role of partnership formation with UK Biobank, but our results suggest that divorce/separation in the past two years (we could not evaluate the role of earlier separations) and partner death are unlikely to fully explain the associations we observe.

The use of genetic variants as instrumental variables is an important strength of this analysis, as it means our results are less likely to be biased due to confounding or reverse causality^18^, both of which are important potential sources of bias in observational studies examining the effects of BMI on social and socioeconomic outcomes. Several of our results from MR analyses differ considerably from our analysis using linear/logistic regression. Notably, associations of BMI with educational outcomes (years of schooling and degree-level education) are far weaker when analysed using MR. This suggests that the observational associations are strongly confounded, and is consistent with there being strong effects of socioeconomic background on BMI but only relatively small effects of BMI on education. It also suggests that much of the effect of BMI on income that we observe may not be driven through education, bur rather reflects later life processes. There was also strong evidence of associations between BMI and all measures of social contact in logistic regression analyses, which were not observed in MR analyses (apart from participation in leisure and social activities in men), again likely reflecting confounding of observational analysis. One of the key assumptions of MR is that the genetic variants used as an instrumental variable affect the outcome (here, SEP, cohabitation and social contact) only through their effect on the exposure (here, BMI), i.e. we assume the absence of horizontal pleiotropy.^27^ We used sun protection use as a negative control outcome, since it is implausible that BMI has a direct effect on sun protection use. We did find evidence of an effect of BMI on sun protection use, but it was weaker than the associations with SEP. This association could arise through the effect of BMI on SEP, which in turn affects sunscreen use, or alternatively it is possible that the BMI genetic risk score exhibits pleiotropy through other health-promoting behaviours, in which case our estimated associations of BMI with SEP and social outcomes would be overestimated. Our sensitivity analyses using two-sample MR approaches provided results that were generally consistent to our main analysis. The greatest degree of consistency was observed for income and deprivation, which are the two continuous outcomes in our analysis and therefore have the greatest statistical power. For binary outcomes, the two-sample results often had low precision.

An important potential source of bias in our analysis is dynastic effects, i.e. the role of parental genotypes in explaining the observed associations.^32^ For example, parental genotype could influence parental behaviours, which in turn affect their child’s BMI and social and socioeconomic outcomes. The data currently available within UK Biobank does not permit exploration of this issue, but as more studies become available with genetic data for both parents and child, it will be important to evaluate whether the effects of BMI on social and socioeconomic outcomes are robust to consideration of parental genotype.

UK Biobank is restricted to participants born between 1938-1971; the social and socioeconomic consequences of BMI may differ in younger generations who have grown up during the obesity epidemic. Furthermore, UK Biobank had only a 5% response rate amongst those invited to participate.^23 33^ It is a relatively homogenous population, and our analyses were restricted to people of white European descent; the generalisability of our findings may therefore be limited. Such patterns of selection have previously been shown to underestimate socioeconomic inequalities in health^34^, and it is possible that the effects of BMI estimated in this study are underestimates of those seen in the broader UK population. Non-linear MR analysis has lower statistical power than linear MR models, and it is therefore possible that for some outcomes where we do not show non-linear effects, these exist but we lack statistical power to detect them.

In the previous analysis of a smaller subset of UK Biobank participants, the effects of BMI on SEP were stronger in women.^19^ In this updated analysis in almost three times the number of participants, the gender differences for effects of BMI on SEP are less pronounced. However, we observe strong gender differences in the effects of BMI on odds of being in a cohabiting relationship. In men, there was strong evidence of low BMI being associated with reduced odds of cohabitation with a partner or spouse. In contrast, for women there was a strong effect of higher BMI on lower odds of cohabitation with a partner or spouse. This finding may be indicative of gender differences in the cultural idealisation and social values for body size, with thinness being culturally valued in women, but perceived strength being valued in men.^35^ Whilst there is evidence that cohabitation can have a protective effect on mortality^36^, there is also evidence of advantages of being single, including more expansive social networks.^37^

In summary, we use Mendelian Randomization to demonstrate effects of both high and low BMI on SEP and odds of being in a cohabiting relationship, with important gender differences in the effects of BMI on cohabitation. Our results have important implications for public health and public policy more broadly. The prevalence of obesity is rising globally^38^ ^39^, and our results suggest this may have adverse socioeconomic consequences for individuals and populations, with consequences likely to accrue across the life course, from lower educational attainment to lower income. Successful obesity prevention and treatment, in addition to health benefits, may also reduce socioeconomic inequalities. Whilst there remain gaps in our understanding, specifically how the relationships differ across populations and generations, and understanding the role of parental genotype, the causal link between BMI and socioeconomic outcomes presented here contributes towards building the case for action to support healthy BMI in this generation and the next.

## Data sharing

The data reported in this paper are available via application directly to UK Biobank. This work was conducted using the UK Biobank under application 9072.

## Ethics approval

UK Biobank has received ethics approval from the National Health Service National Research Ethics Service (ref 11/NW/0382).

## Transparency statement

LDH and JT (study guarantors) affirm that the manuscript is an honest, accurate, and transparent account of the study being reported; that no important aspects of the study have been omitted; and that any discrepancies from the study as originally planned have been explained.

