## Supplementary Text for "Effects of body mass index on relationship status, social contact, and socioeconomic position: Mendelian Randomization study in UK Biobank"

Laura D Howe, reader*^1^; Roshni Kanayalal, research intern^2^; Robin N Beaumont, research fellow^2^; Alisha R Davies, head of research and development^3^; Tim Frayling^2^, Sean Harrison, senior research associate^1^; Samuel E Jones, research fellow^2^; Franco Sassi, professor^4^; Andrew R Wood, research fellow^2^; Jessica Tyrrell, lecturer^2^

1 – MRC Integrative Epidemiology Unit at the University of Bristol, Population Health Sciences, Bristol Medical School, University of Bristol, Bristol, BS8 2BN, UK

2 – Genetics of Complex Traits, University of Exeter Medical School, RILD Level 3, Royal Devon & Exeter Hospital, Barrack Road, Exeter EX2 5DW, UK

3 – Public Health Wales, 2 Capital Quarter, Tyndall St, Cardiff CF10 4BZ

4 – Centre for Health Economics and Policy Innovation, Imperial College Business School, London, UK.

**Supplementary Material**

**UK Biobank variables used**

These analyses used the following UK Biobank variables as exposures and outcomes:

- n_21001_0_0: Body mass index
- n_189_0_0: Townsend deprivation index
- n_738_0_0: Annual household income
- n_132_0_0: Job code
- n_845_0_0: Age completed full time education
- n_6138_0_0: Qualifications
- n_6142_0_0: Current employment status
- n_1031_0_0: Visits from friends and family
- n_6160_0_0: Leisure/social activities
- n_2110_0_0: Able to confide
- n_6141_0_0: How are people in household related to participant

**Details of socioeconomic position measures used**

- Townsend deprivation index (TDI); an area-based measure at census-tract level calculated by the indicators: non-home ownership, non-car ownership, unemployment and overcrowded households. A continuous variable; higher scores indicate higher deprivation.
- Annual household income, a questionnaire-reported variable with 5 categories: <£18,000, £18,000 to £30,999, £31,000 to £51,999, £52,000 to £100,000, and >£100,000. This was not available for participants who indicated they were living in sheltered accommodation or in a care home. This variable was an ordinal scale. To convert to estimates of income the midpoint of each group was utilised.
- Job class; coded as managers and senior officials, professional occupations, associate professionals, business and public sector associate professionals, admin and secretarial roles, skills trade, personal service occupations, leisure and other personal service occupations, sales and customer. We created a dichotomous variable where higher skilled represented managers and senior officials to skills trade and lower skilled the remaining 5 categories.
- Employment status, based on participants’ responses to “Which of the following describes your current situation”, with the options to respond: a) In paid employment or self-employed, b) Retired, c) Looking after home and/or family, d) Unable to work because of sickness or disability, e) Unemployed, f) Doing unpaid or voluntary work, g) Full or part-time student, h) None of the above. We created a binary variable comparing participants reporting that they were solely unemployed to those reporting they were employed/self-employed; other responses were coded as missing.
- Years in education was derived from two questions in the UK Biobank. Firstly, whether participants had a college or university degree. If they did not have a degree, they were asked what age they left full-time education. Participants with a degree were coded as having completed full-time education at 21 years of age.
- Degree status; participants reported their qualifications in a questionnaire. We created a dichotomous variable for degree level and professional (e.g. nursing or teaching) qualifications versus other qualifications; participants responding ‘prefer not to answer’ were coded as missing.

**Details of social contact measures**

- Visits from friends and family: participants were asked how frequently they saw friends and family with the option to report never or almost never, once every few months, about once a month, about once a week, 2-4 times a week, almost daily. We derived a dichotomous variable; less than weekly versus weekly or more visits.
- Participation in leisure and social activity: participants were asked "Which of the following do you attend once a week or more often? (You can select more than one)", with the option to report sports club or gym, social club, religious group, educational class, other group activity or none of the above. We created a dichotomous variable of any activity versus none.
- Confiding in others: participants were asked “How often are you able to confide in someone close to you?”. We created a dichotomous variable of less than weekly versus weekly or more.

**Details of genotyping, genetic QC, sample definition, and relatedness**

Genotypes were generated from the Affymetrix Axiom UK Biobank array (~450,000 individuals) and the UKBiLEVE array (~50,000 individuals). This dataset underwent extensive central quality control (<http://biobank.ctsu.ox.ac.uk)>. Principal components were generated in the 1000 Genomes Cohort using high-confidence SNPs. These were SNPs that overlapped with SNPs directly genotyped in the UK Biobank, with a MAF>5% and HWE>1x10^-6^ in the 1000 Genomes cohort. In the UK Biobank, SNPs had to pass the UK Biobank defined quality control across all batches and missingness <1.5%.^30^ Principal component analysis was then used to obtain their individual loadings. These loadings were then used to project all of the UK Biobank samples into the same principal component space and individuals were then clustered using principal components 1 to 4. Participants were removed if they had subsequently withdrawn from the study (n=7), if they were sex mismatches (n=348; self-reported sex did not match genetic sex), or, if they were related up to third degree to another participant in the cohort (n=71,123). Related individuals were defined using a KING Kinship and an optimal list of unrelated individuals was generated to allow maximum numbers of individuals to be included. Ancestral principal components were then generated within these identified individuals for use in subsequent analyses.

**Details of statistical methods for two sample MR methods to assess pleiotropy**

Each BMI variant was regressed against each outcome using the appropriate model (linear or logistic regression) and the beta coefficient and standard error extracted. We performed Inverse Variance Weighted (IVW) instrumental variable analysis and two methods that are more robust to potential violations of the standard instrumental variable assumptions (MR-Egger^1^ and Median MR^2^). The two-sample approach regresses the effect sizes of variant-outcome associations (here BMI variants versus SEP, cohabitation and social contact outcomes) against effect sizes of the variant-risk factor associations (here BMI variants versus BMI, from the primary GWAS of BMI^3^).

The IVW approach assumes no horizontal pleiotropy (under a fixed effect model) or, if implemented under a random effects model after detecting heterogeneity amongst the causal estimates, that: a) the strength of the association of the genetic instruments with the risk factor is not correlated with the size of the pleiotropic effects and b) the pleiotropic effects have an average value of zero.

In contrast, the MR-Egger performs weighted regression with an unconstrained intercept. This removes the assumption that all genetic variants are valid instruments and is therefore less susceptible to confounding from pleiotropic variants, that have a stronger effect on the outcome than the primary trait. Median-MR is robust when up to 50% of the genetic variants are invalid. It takes the median instrumental variable from all variants included. If all the methods are broadly consistent it strengthens our causal inference. Details of the R code for the 2-sample IVW, Egger and Median analyses are provided in Bowden et al., 2015 and 2016.^1 2^

**Details of statistical methods for non-linear MR**

The nlmr package in R regresses the exposure (here, BMI) on the instrumental variable (genetic risk score for BMI) to generate the ‘IV-free’ exposure (non-genetic component of BMI). In strata of the IV-free exposure, the local average causal effect (LACE) of BMI on the outcome is estimated as a ratio of coefficients: the IV association with the outcome divided by the IV association with the exposure. This approach assumes a linear effect of the IV on the exposure. The nlmr package provides two options for estimating the non-linear effects of an exposure on an outcome; fractional polynomials and a piecewise linear function. Fractional polynomial methods can be unduly influenced by the extremes of a distribution, therefore we used the piecewise linear function only. The piecewise linear method estimates a continuous function, whereby a linear relationship is fitted within each stratum of the IV-free exposure distribution, constrained so that each segment begins where the previous one ended. Confidence intervals are estimated by bootstrapping. We a priori selected to run our analysis across deciles of IV-free BMI. Two statistical tests of non-linearity are presented: Cochran’s Q statistic assesses whether heterogeneity of LACE estimates is greater than would be expected by chance, and a quadratic test metaregresses the LACE estimates against the mean exposure value in each stratum (equivalent to fitting a quadratic exposure-outcome model).

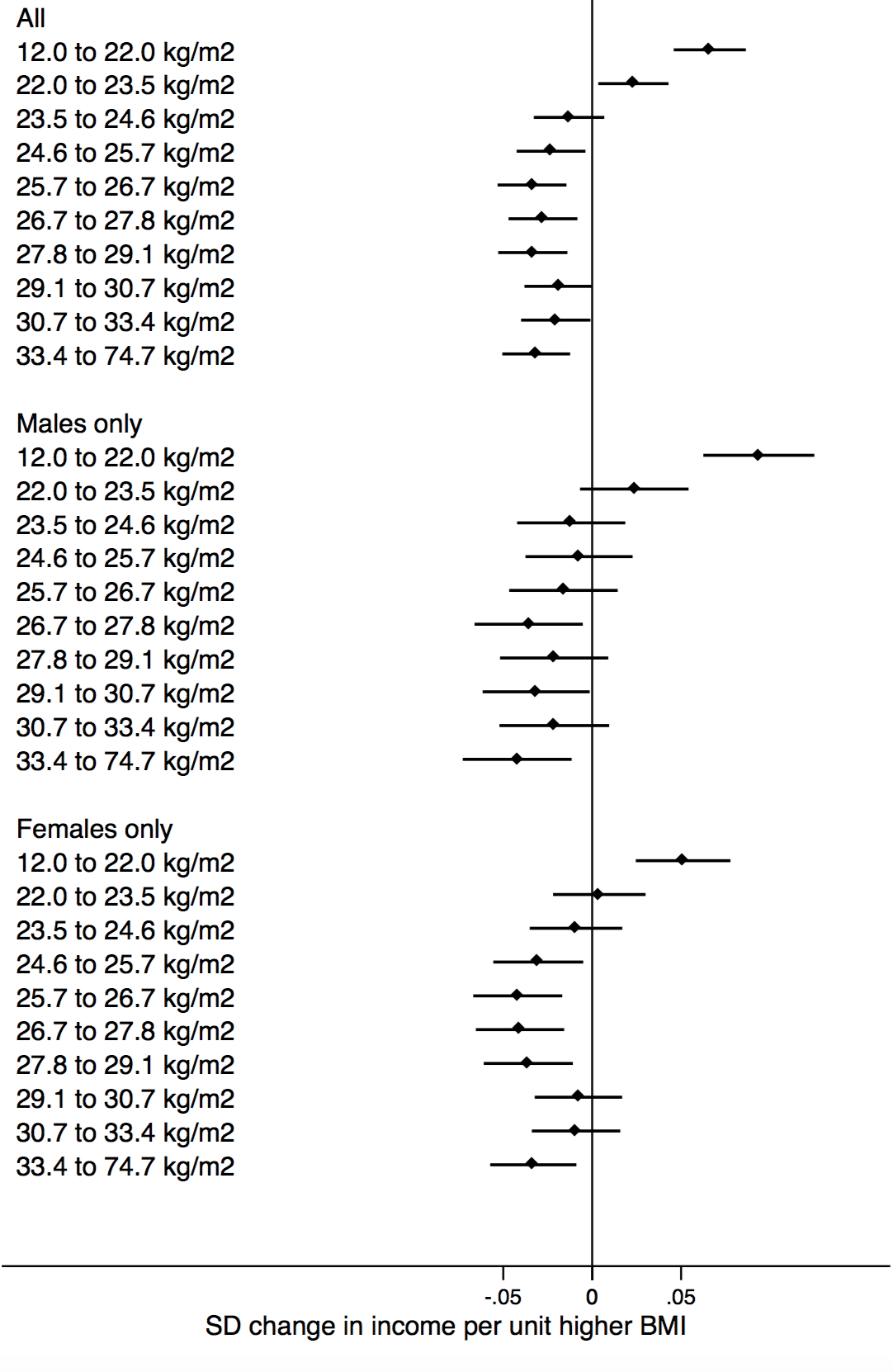

P_Quadratic_: 4x10^-9^

P_Cochran Q_: 5x10^-15^

P_Quadratic_: 2x10^-9^

P_Cochran Q_: 5x10^-9^

P_Quadratic_: 0.004

P_Cochran Q_: 5x10^-6^

Lower income

Higher income

**Supplementary figure 1:** Dot plot exploring the non-linear association between BMI and income in 10 BMI strata using the Piecewise Linear Model from non-linear MR. The p-values presented for causal non-linear relationships are from the Piecewise Linear Model (P_Quadratic_ and P_Cochran Q_). Full details of non-linear analyses in Supplementary table 6.
